# Myosin Light Chain Dephosphorylation by PPP1R12C Promotes Atrial Hypocontractility in Atrial Fibrillation

**DOI:** 10.1101/2023.04.19.537590

**Authors:** Srikanth Perike, Francisco J. Gonzalez-Gonzalez, Issam Abu-Taha, Frederick W. Damen, Ken S. Lizama, Anahita Aboonabi, Andrielle E. Capote, Yuriana Aguilar-Sanchez, Benjamin Levin, Zhenbo Han, Arvind Sridhar, Jacob Grand, Jody Martin, Joseph G. Akar, Chad M. Warren, R. John Solaro, Sang-Ging Ong, Dawood Darbar, Craig J. Goergen, Beata M. Wolska, Dobromir Dobrev, Xander H.T. Wehrens, Mark D. McCauley

**Affiliations:** Division of Cardiology, Department of Medicine, College of Medicine, University of Illinois at Chicago, Chicago, Illinois, USA; Jesse Brown VA Medical Center, Chicago, Illinois, USA; Department of Physiology and Biophysics and the Center for Cardiovascular Research, and the College of Medicine, University of Illinois at Chicago, Chicago, Illinois, USA; Department of Pharmacology and Regenerative Medicine, College of Medicine, University of Illinois at Chicago, Chicago, Illinois, USA; Institute of Pharmacology, West German Heart and Vascular Center, University Duisburg-Essen, Essen, Germany; Weldon School of Biomedical Engineering, Purdue University, West Lafayette, IN, USA; Department of Integrative Physiology and The Cardiovascular Research Institute, Baylor College of Medicine, Houston, TX, USA; Oregon Health and Science University, Portland, OR, USA; University of California Davis, Davis, CA, USA; Yale University, New Haven, CT, USA; Department of Medicine, Montréal Heart Institute and Université de Montréal, Montréal, Canada

**Keywords:** Atrial fibrillation, myosin light chain, cardiac arrhythmias, protein phosphatase 1, protein phosphatase regulatory subunit 12C

## Abstract

**Background:** Atrial fibrillation (AF), the most common sustained cardiac arrhythmia, increases thromboembolic stroke risk five-fold. Although atrial hypocontractility contributes to stroke risk in AF, the molecular mechanisms reducing myofilament contractile function remain unknown. We tested the hypothesis that increased expression of PPP1R12C, the PP1 regulatory subunit targeting atrial myosin light chain 2 (MLC2a), causes hypophosphorylation of MLC2a and results in atrial hypocontractility.

**Methods:** Right atrial appendage tissues were isolated from human AF patients versus sinus rhythm (SR) controls. Western blots, co-immunoprecipitation, and phosphorylation studies were performed to examine how the PP1c-PPP1R12C interaction causes MLC2a de-phosphorylation. *In vitro* studies of pharmacologic MRCK inhibitor (BDP5290) in atrial HL-1 cells were performed to evaluate PP1 holoenzyme activity on MLC2a. Cardiac-specific lentiviral PPP1R12C overexpression was performed in mice to evaluate atrial remodeling with atrial cell shortening assays, echocardiography, and AF inducibility with EP studies.

**Results:** In human patients with AF, PPP1R12C expression was increased two-fold versus SR controls (*P*=2.0×10^−2^, n=12,12 in each group) with > 40% reduction in MLC2a phosphorylation (*P*=1.4×10^−6^, n=12,12 in each group). PPP1R12C-PP1c binding and PPP1R12C-MLC2a binding were significantly increased in AF (*P*=2.9×10^−2^ and 6.7×10^−3^ respectively, n=8,8 in each group). *In vitro* studies utilizing drug BDP5290, which inhibits T560-PPP1R12C phosphorylation, demonstrated increased PPP1R12C binding with both PP1c and MLC2a, and dephosphorylation of MLC2a. Lenti-12C mice demonstrated a 150% increase in LA size versus controls (*P*=5.0×10^−6^, n=12,8,12), with reduced atrial strain and atrial ejection fraction. Pacing-induced AF in Lenti-12C mice was significantly higher than controls (*P*=1.8×10^−2^ and 4.1×10^−2^ respectively, n= 6,6,5).

**Conclusions:** AF patients exhibit increased levels of PPP1R12C protein compared to controls. PPP1R12C overexpression in mice increases PP1c targeting to MLC2a and causes MLC2a dephosphorylation, which reduces atrial contractility and increases AF inducibility. These findings suggest that PP1 regulation of sarcomere function at MLC2a is a key determinant of atrial contractility in AF.

## INTRODUCTION

Atrial fibrillation (AF) is the most common sustained cardiac arrhythmia, and is an independent risk factor for the development of stroke, thromboembolism, heart failure, and impaired quality of life.^1-3^ Stroke remains one of the most dreaded and deadly complications of AF. Recent studies have shown that even subclinical AF more than doubles a patient’s risk for stroke in as little as three months^4,5^, and up to 40% of cryptogenic strokes are attributable to AF.^6^ The mechanisms of thrombogenesis in AF are multifactorial, with evidence to suggest that AF is associated with all three criteria of Virchow’s triad – abnormal blood stasis, endothelial damage, and hypercoagulability. Current therapies to reduce stroke risk in AF include anticoagulants and left atrial appendage (LAA) exclusion, which address two of the three factors of Virchow’s triad, namely hypercoagulability of blood and exclusion of the most thrombogenic portion of the left atrium, the left atrial appendage.^7–10^ However, to date, there are no therapeutic agents that directly reverse atrial cardiomyopathy or improve atrial contractility. Furthermore, recent mechanistic insights have led to the recognition that atrial hypocontractility is a key contributor to atrial cardiomyopathy and is a substrate for AF.^11^ Given the importance of atrial cardiomyopathy in stroke risk, we sought to examine the undSerlying mechanisms of atrial hypocontractility, or atrial stunning, that may serve as a substrate for AF.

The cellular mechanisms responsible for atrial hypocontractilty in AF remain poorly understood.^12^ Shortening of the atrial effective refractory period (AERP), intracellular Ca^2+^ overload, changes in myofilament Ca^2+^ sensitivity, mitochondrial dysfunction, and pro-apoptotic and pro-fibrotic signaling have all been described in AF.^13–16^ Central to the mechanism of AF-induced hypocontractility is dysregulation of atrial myofilament signaling. Cardiac contractility relies upon phosphorylation processes, which in AF are pathologically altered.^13^ For example, in human patients with chronic AF, there is an increase in total protein phosphatase 1 (PP1) activity, which is associated with dephosphorylation of key regulatory proteins involved in Ca^2+^ homeostasis.^14, 16^ More recently, Chiang et al.^17^ performed a detailed PP1 interactome analysis in human paroxysmal AF, and determined that alterations in PP1 regulatory (PPP1R) subunit binding to the catalytic subunit (PP1c) may underlie pathologic signaling for atrial excitation-contraction coupling. Of these interactions, only one regulatory subunit, PPP1R12C, has been identified that targets the sarcomere at a key regulatory site for atrial contractility, atrial myosin light chain 2 (MLC2a).^18, 19^ However, the interactome study was underpowered to determine whether PPP1R12C is significantly increased in human AF. Also, further knowledge gaps remained as to whether overexpression of PPP1R12C in human AF is sufficient to dephosphorylate MLC2a and drive atrial remodeling leading to atrial hypocontractility and AF risk.

Given the importance of atrial hypocontractility in stroke risk, evidence of enhanced PP1 activity in AF, and a recent link between PP1c-PPP1R12C in AF, we tested the hypothesis that increased expression of PPP1R12C, the PP1 regulatory subunit targeting atrial myosin light chain 2 (MLC2a), causes hypophosphorylation of MLC2a, which in turn results in atrial hypocontractility and susceptibility to pacing-induced AF.

## METHODS

### Human Atrial Samples

All patients participating in this study gave written informed consent according to the Declaration of Helsinki, and the study was approved by the institutional review committees. Right atrial appendages were dissected from patients undergoing open heart surgery, and tissues were flash-frozen in liquid nitrogen. Experimental protocols were approved by the ethics committee at the University Duisburg-Essen (#12-5268-BO) and the University of Illinois at Chicago (2015-1149).

### Experimental Mice

All animal studies were performed according to protocols approved by the Animal Care and Use Committee (ACC#20-005) at UIC conforming to the *Guide for the Care and Use of Laboratory Animals* published by the U.S. National Institutes of Health. Whole atria were harvested from male and female mice with a C57BL/6 background between the ages of 2-4 months and were immediately flash-frozen in liquid nitrogen. Groups include wild-type (WT) mice, mice treated with lentiviral GFP control (Lenti-Ctl), and mice treated with lentivirus with PPP1R12C-GFP gene insert (Lenti- 12C).

### Western Blot Analysis

50 μg of mouse or human atrial sample lysates were subjected to electrophoresis on 4-12% acrylamide gels, and were transferred onto polyvinyl difluoride (PVDF) membranes. Antibodies against the following targets were used to probe the membranes: rabbit polyclonal anti-PP1c (1:1000, Abcam, Cambridge, MA), rabbit polyclonal anti-PPP1R12C (1:1000, Thermofisher, Rockford, IL), rabbit polyclonal anti-MLC2a (1:2000, Proteintech, Rosemont, IL), and rabbit polyclonal anti-P-MLC2a (1:1000, Cell Signaling, Danvers, MA). Membranes were then incubated with corresponding secondary anti-goat, anti-mouse, anti-chicken and anti-rabbit antibodies that are conjugated to horseradish peroxidase (1: 2000, Abcam, Cambridge, MA), respectively. Linearity of PPP1R12C antibody signal was determined by Western Blot (**Supplemental Figure 1**). Bands were quantified using ImageJ software.^20^ PPP1R12C and MLC2a proteins were normalized vs. β-actin protein, as were PP1c and MLCK proteins. Details of phosphorylation assays, co-immunoprecipitation, and RT-PCR methods are included in the Online Supplement.

### Cell Culture and MRCK inhibition

HL-1 cells were obtained from Dr. Wehrens’ laboratory and were initially purchased from Millipore-Sigma. Passage number ranged from 7-10. HL-1 cells were cultured in Claycomb’s medium (Sigma Aldrich, St. Louis, MO) supplemented with 10% fetal calf serum (FCS), 100 µg/ml Pencillin/Streptomycin, 0.1 mM epinephrine and 2 mM L-Glutamine. Cultures were maintained at 37°C in humidified air and 5% CO_2_. Cells were detached by trypsinization and its activity was neutralized by Soybean trypsin inhibitor (Life technologies, Grand Island, NY). Culture conditions for HL-1 cells have been described previously.^21^ HL-1 cells were incubated with a MRCK inhibitor for 30 min at 37°C (BDP5290, Aobious, Gloucester, MA). The MRCK inhibitor was solubilized in DMSO and the final concentration of DMSO in the culture medium never exceeded 0.1%. DMSO treated cells were used as controls. The dose response of MRCK inhibitor in MLC dephosphorylation have been described.^22^ **Lentivirus-mediated cardiac specific PPP1R12C expression in neonatal mouse hearts *in vivo***. Lentiviruses encoding cardiac specific PPP1R12C and lentivirus control containing 10 µL of 10^9^ particles were injected directly into neonatal mouse hearts from days 1-5 after birth. Mice were sacrificed after 6 weeks post-lentiviral infections for protein expression analysis using Western blot analyses. See Supplemental Methods.

### Cell Shortening Assays

Langendorff perfusion technique was used to perform atrial isolation. 260 μl of the resultant cell suspension was loaded into an imaging chamber with field stimulation capability (RC-21BRFS, Warner Instruments, Connecticut). Atrial cells were stimulated with pacing at 1 Hz frequency at 15 Volts for 3 seconds. Video microscopy was acquired using a Zeiss Laser TIRF spinning disk confocal microscope with a resolution of 90 frames per second. Video analysis was performed using Muscle- Motion open-source software to obtain all reported parameters.

### Transmission Electron Microscopy

Specimens were fixed in 2.5% glutaraldehyde (pH 7.2), post-fixed with 1% osmium tetroxide (1 hr) and dehydrated using an ascending series of ethanol (through 100% absolute). Specimens were then embedded in LX112 epoxy resin and polymerized at 60C for 3 days. Thin sections (∼75 nm) were collected onto copper grids and stained with uranyl acetate and lead citrate, respectively. Specimens were examined using a JEOL JEM-1400F transmission electron microscope (at 80 kV). Micrographs were acquired using an AMT Side-Mount Nano Sprint Model 1200S-B camera, loaded with AMT Imaging software V.7.0.1.

### Echocardiography

Echocardiographic images were acquired using a Vevo2100 high frequency ultrasound system with a 22-55MHz frequency transducer (MS550D; FUJIFILM VisualSonics Inc., Toronto, Canada). Measurements were made while the mouse was anesthetized using 1-2% isoflurane and heart rate > 400 bpm. Body temperature was maintained at 37±1°C. A detailed discussion of atrial measurements^23, 24^ and strain analysis^25, 26^ is included in the Online Supplement.

### Programmed Electrical Stimulation

Trans-esophageal (TE) pacing of the atria was performed using a 1.1F octapolar catheter (Millar Instruments, Houston, TX)^20^ inserted into the esophagus to the level of the heart, as described in the Online Supplement.^27, 28^

### Statistical Analysis

Statistical analysis was performed using Graphpad Prism 9.0.1. Data are expressed as means ± standard deviation (SD). Numeric data were first analyzed for normality using the Shapiro-Wilk test. Data with a parametric distribution were analyzed by an unpaired two-tailed Student’s *t*-test and one-way ANOVA. When significant departures from normality were observed by the Shapiro-Wilk test, or when datasets were too small to robustly test whether the assumptions of parametric testing were met, non-parametric tests were used. For echocardiograms, one-way ANOVA and Tukey’s post-hoc multiple comparisons test were used. Representative images and figures were chosen based on their proximity to the mean/average for each group.

## RESULTS

### PPP1R12C Expression is Increased in Human AF

To confirm the role of PPP1R12C in AF, we first used human atrial appendage samples from de-identified patients with AF versus sinus rhythm (SR). Western blotting of atrial lysates showed a two-fold increase of PPP1R12C protein expression in AF samples versus SR controls (n = 12,12 in each group; *P* = 2.0×10^−2^; Figure 1A and 1B), as well as a significant increase in *PPP1R12C* gene transcription (n = 5,5 each; *P* = 1.5×10^−2^; **Supplemental Figure 2**). Increased PPP1R12C expression was associated with a significant reduction in MLC2a phosphorylation in AF versus SR (n = 12,12 in each group; *P* = 1.4×10^−6^, Figure 1C-D), without a significant change in MLC2a expression or catalytic PP1c expression (n = 12,12 in each group; *P* = 3.5×10^−1^; Figure 1E-1F). However, Western blotting of myosin light chain kinase (MLCK), the predominant kinase regulating MLC2a phosphorylation, showed no significant changes in expression (**Supplemental Figure 3A-B**), suggesting that increased expression of PPP1R12C and increased targeting of PP1 to the sarcomere could be the major determinant of MLC2a dephosphorylation in human AF. **Supplemental Table 1** shows a comparison of human patient clinical data between the two groups.

**Figure 1.**
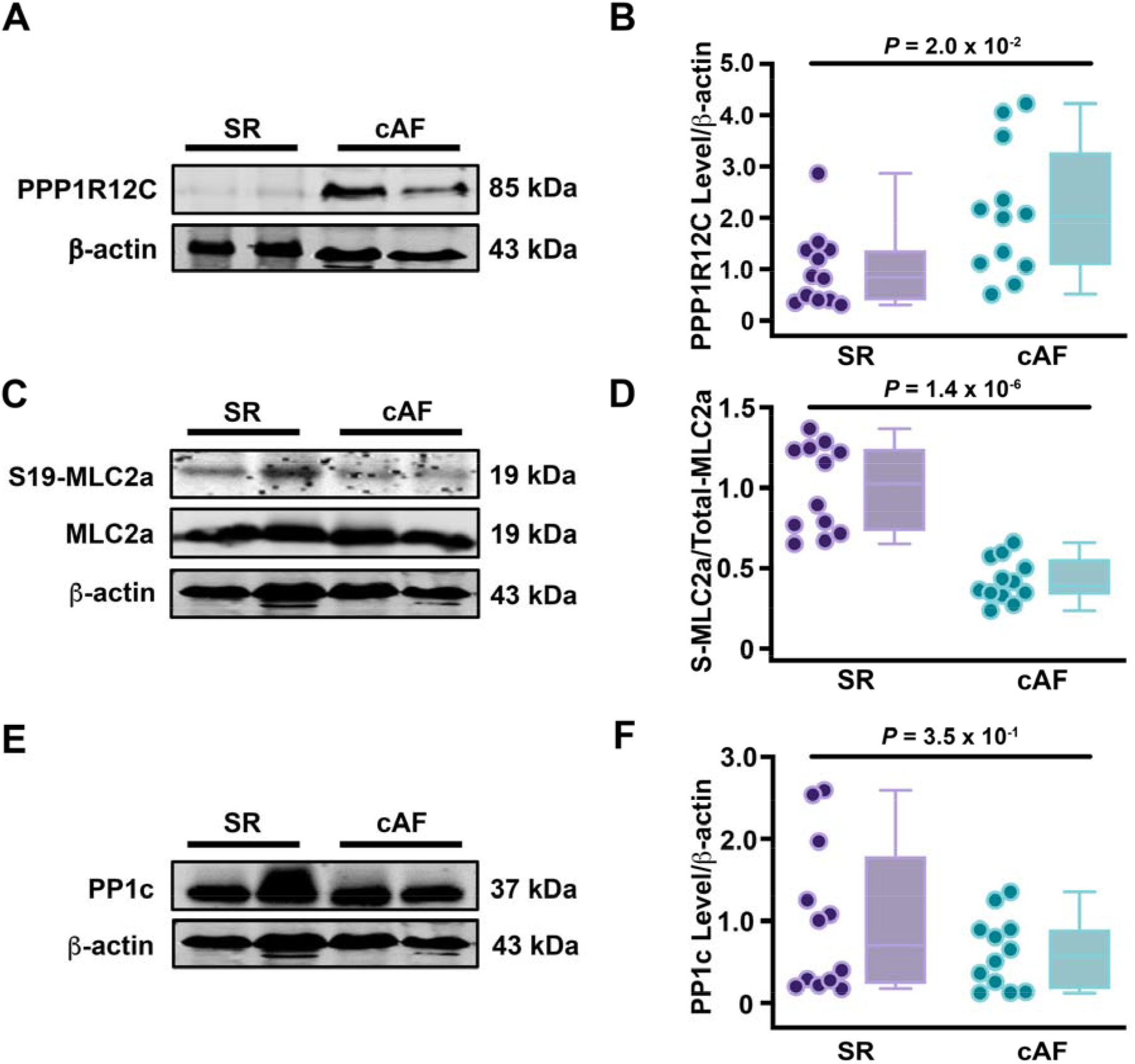
Increased PPP1R12C expression in human AF is associated with reduced MLC2a phosphorylation. **A**, Representative Western blot comparing the expression of PPP1R12C in human atrial tissues with SR or cAF. **B**, Quantification of PPP1R12C protein abundance. n=12 in each group. **C**, Representative Western blot and phosphorylation assays comparing the phosphorylation and abundance of MLC2a in cAF and SR tissues. **D**, Quantification of fold change in phosphorylation of MLC2a. n=12 in each group. **E**, Western blot comparing the expression of PP1c in human atrial tissues with SR or cAF. **F**, Quantification of PP1c abundance in AF versus SR atrial tissues. n=12 in each group. Data represent mean±SD. Data were determined to have a non-parametric distribution by the Shapiro-Wilk test, and were analyzed using the Mann-Whitney test. AU indicates arbitrary units; cAF, chronic atrial fibrillation; MLC2a, myosin light chain 2a; PP1c, protein phosphatase 1 catalytic subunit; PPP1R12C, protein phosphatase 1 regulatory subunit 12C; SR, sinus rhythm.

We next evaluated the protein binding relationships among the regulatory targeting protein PPP1R12C, the PP1 catalytic domain (PP1c), and MLC2a in human AF and SR samples. Co-immunoprecipitation (co-IP) demonstrated an enhanced interaction between PP1c and PPP1R12C in human patients with AF versus SR patients (n = 8,8 in each group; *P* = 2.9×10^−2^; Figure 2A and 2B, **Supplemental Table 2**). Likewise, co-IP showed a stronger interaction between MLC2a and PPP1R12C in AF versus SR (n = 8,8 in each group; *P* = 6.7×10^−3^; Figure 2C and 2D). These results are consistent with prior reports in neuronal tissue that PPP1R12C (also called MBS85) serves as a scaffold upon which PP1c dephosphorylates MLC2, and provide a mechanistic basis for the de-phosphorylation of MLC2a in AF.^19^

**Figure 2.**
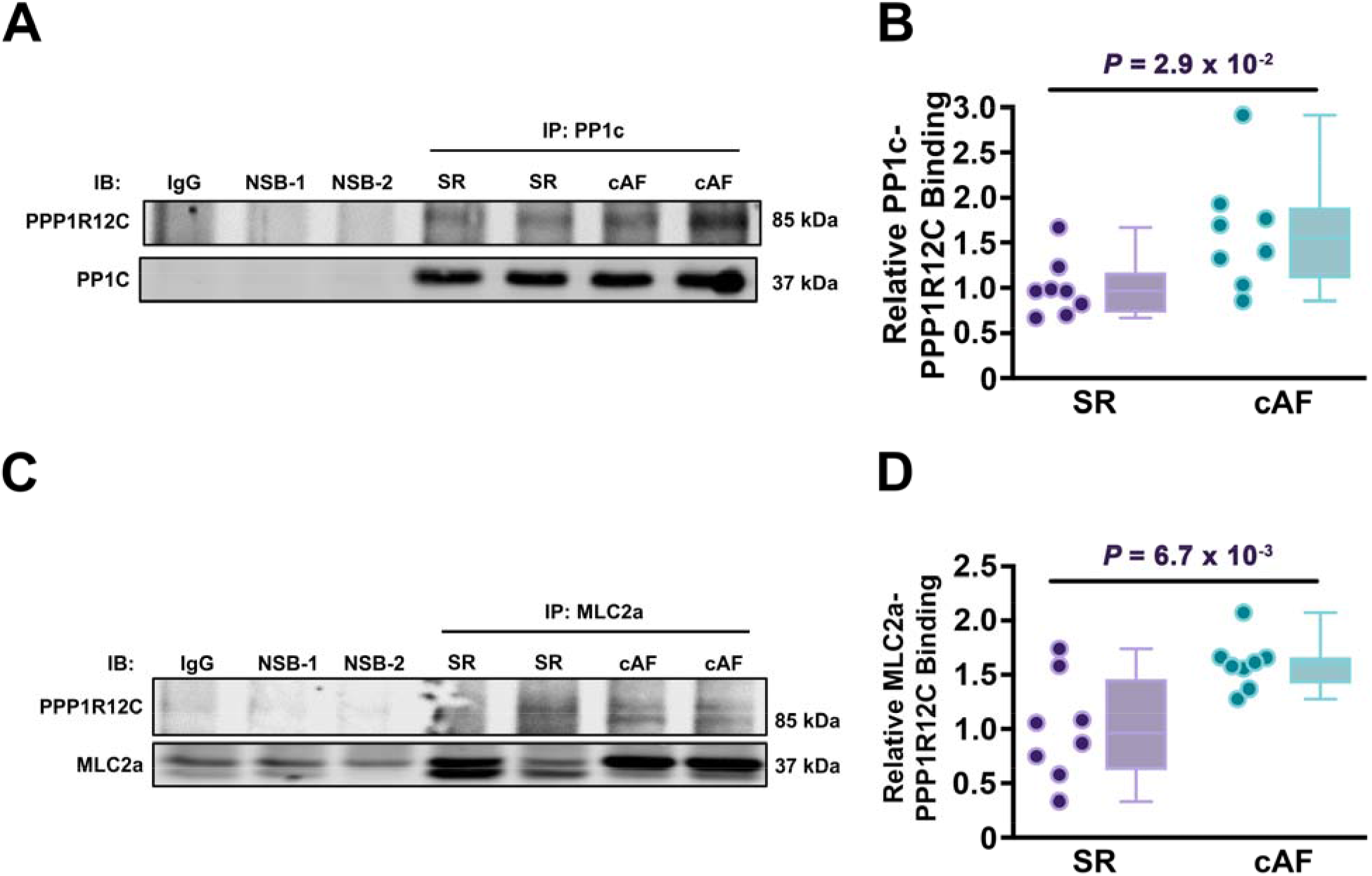
Enhanced PPP1R12C binding to PP1c and MLC2a in cAF versus SR. **A**, Representative co-immunoprecipitation assay comparing PPP1R12C and PP1c binding in human cAF and SR atrial tissues. **B**, Quantification of PPP1R12C-PP1c protein binding. **C**, Representative co-immunoprecipitation assay showing PPP1R12C-MLC2a binding in human cAF and SR. **D**, Quantification of PPP1R12C-MLC2a binding. Data represent mean±SD. Data were determined to have a parametric distribution by the Shapiro-Wilk test, and were analyzed using unpaired 2-tailed Student’s *t*-test. cAF indicates chronic atrial fibrillation; IB, immunoblot; IgG, immunoglobulin G antibody control; IP, immunoprecipitate; MLC2a, myosin light chain 2a; NSB, non-specific binding negative control; PP1c, protein phosphatase 1 catalytic subunit; PPP1R12C, protein phosphatase 1 regulatory subunit 12C; SR, sinus rhythm.

### PPP1R12C Activity is Modifiable by RhoA Pathway Signaling

Tan *et al*. previously identified T560-PPP1R12C as a key phosphorylation site on PPP1R12C protein, which alters PPP1R12C binding activity, and is phosphorylated by myotonic dystrophy kinase-related Cdc-42 binding kinase (MRCK) via the ras homolog family A (RhoA) signaling pathway.^19^ Given the importance of RhoA pathway signaling in promoting pro-inflammatory and pro-fibrotic remodeling in AF^29–31^, we sought to inhibit phosphorylation at this regulatory site to determine if we could modify binding of PP1c- PPP1R12C and PPP1R12C-MLC2a *in vitro*. We utilized a pharmacologic inhibitor of MRCK (BDP5290) to determine if the T560-PPP1R12C site could be a potential drug target for future modification of PPP1R12C activity and MLC2a phosphorylation. As predicted, BDP5290 activated PPP1R12C, enhanced binding to PP1c (n = 8,8,9; *P* = 1.3×10^−2^; Figure 3A-B) and MLC2a (n = 6 in each group; *P* = 6.0×10^−4^; Figure 3C-D), and contributed to MLC2a dephosphorylation (n = 5 in each group; *P* = 4.1×10^−2^; Figure 3E-F). These results demonstrate that rather than a passive molecular scaffold, PPP1R12C is a dynamic molecule that responds to Rho/ROCK signaling pathways via MRCK, can modulate MLC2a phosphorylation, and can alter PP1c binding to MLC2a.

**Figure 3.**
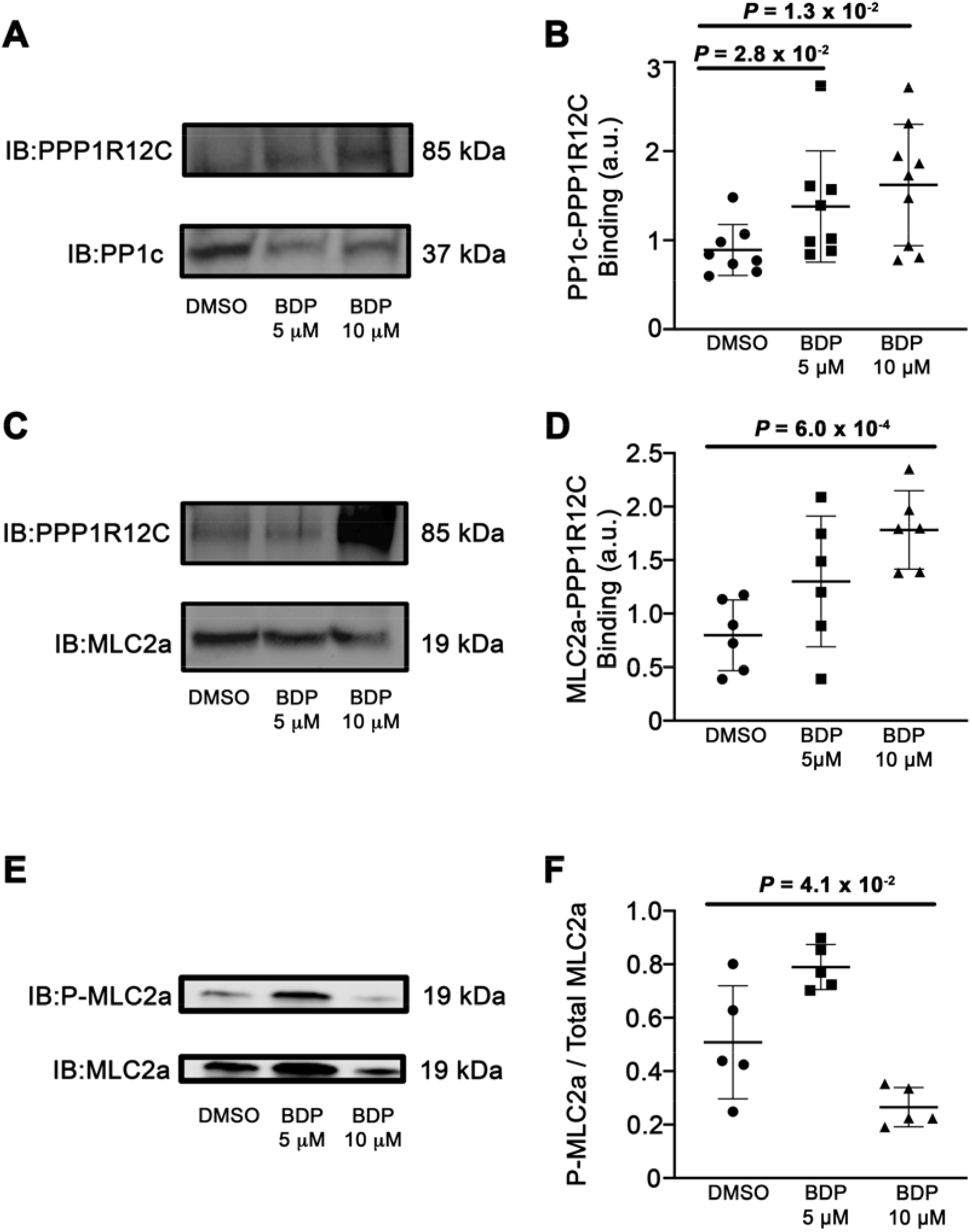
Dephosphorylation of T560-PPP1R12C promotes binding to PP1c and MLC2a. **A**, Co-IP studies demonstrating how the MRCK-inhibiting drug BDP5290 increases binding between PPP1R12C and PP1c versus DMSO control. **B**, Quantification of protein binding between PPP1R12C and PP1c in HL-1 cells. n = 8,8,9 in each group. **C**, Co-IP studies demonstrating that BDP5290 increases binding between PPP1R12C and MLC2a in atrial HL-1 cells. **D**, Quantification of protein binding between PPP1R12C and MLC2a. n = 6 per group. **E**, Phosphorylation assay showing the effects of BDP5290 on MLC2a phosphorylation in treated atrial HL-1 cells. **F**, Quantification of phosphorylation normalized to DMSO negative controls. n = 5 per group. Data represent mean±SD. A Shapiro-Wilk test determined that data distribution is non-parametric for Fig. **3A-B**, and a Mann-Whitney and Kruskal-Wallis test were performed. Shapiro-Wilk tests for Fig. 3C-F showed a parametric distribution, and comparisons were analyzed using an unpaired 2-tailed Student’s *t*-test and ANOVA. AU indicates arbitrary unit; BDP, drug BDP5290; DMSO, dimethylsulfoxide; MLC2a, myosin light chain 2a; P, co-IP pulldown lane; PP1c, protein phosphatase 1 catalytic subunit; PPP1R12C, protein phosphatase 1 regulatory subunit 12C.

### Mice Overexpressing PPP1R12C Develop Atrial Myopathy and Are Prone to Pacing-Induced Atrial Fibrillation

We next sought to determine the functional consequences of altering cardiac PPP1R12C expression upon atrial contractility *in vivo*. To model PPP1R12C overexpression in AF, we established a cardiac-specific lentivirus-mediated PPP1R12C transgenic mouse (Lenti-12C) utilizing both an α-MHC promoter and direct cardiac injection on days 1-5 of postnatal life, according to validated viral vector protocols.^32, 33^ A lentiviral negative control with GFP tag (Lenti-Ctl) was established to differentiate alterations in phenotype from PPP1R12C overexpression versus the use of the lentiviral vector alone. Using this method, we were able to achieve overexpression of PPP1R12C protein in mouse atria at 2-3 months of age versus non-transfected controls (n = 3,4,7 in each group; *P* = 2.8×10^−2^ vs. WT and *P* = 6.0×10^−3^ vs. Lenti-Ctl; Figure 4A-B), which was similar to the level of PPP1R12C expression in human AF samples (Figure 1A-B). Viral transduction efficiency in the heart from this method was > 95% as measured by lenti-GFP labeling (**Supplemental Figure 4**). We measured MLC2a phosphorylation in the three mouse models (WT, Lenti-Ctl, and Lenti-12C). We found that there was a significant decrease in MLC2a phosphorylation in Lenti-12C mice versus WT controls (n = 9,5,8 in each group; *P* = 6.0×10^−4^, Figure 4C); however, there were no significant differences in MLC2a phosphorylation between WT and Lenti-Ctl, as expected.

**Figure 4.**
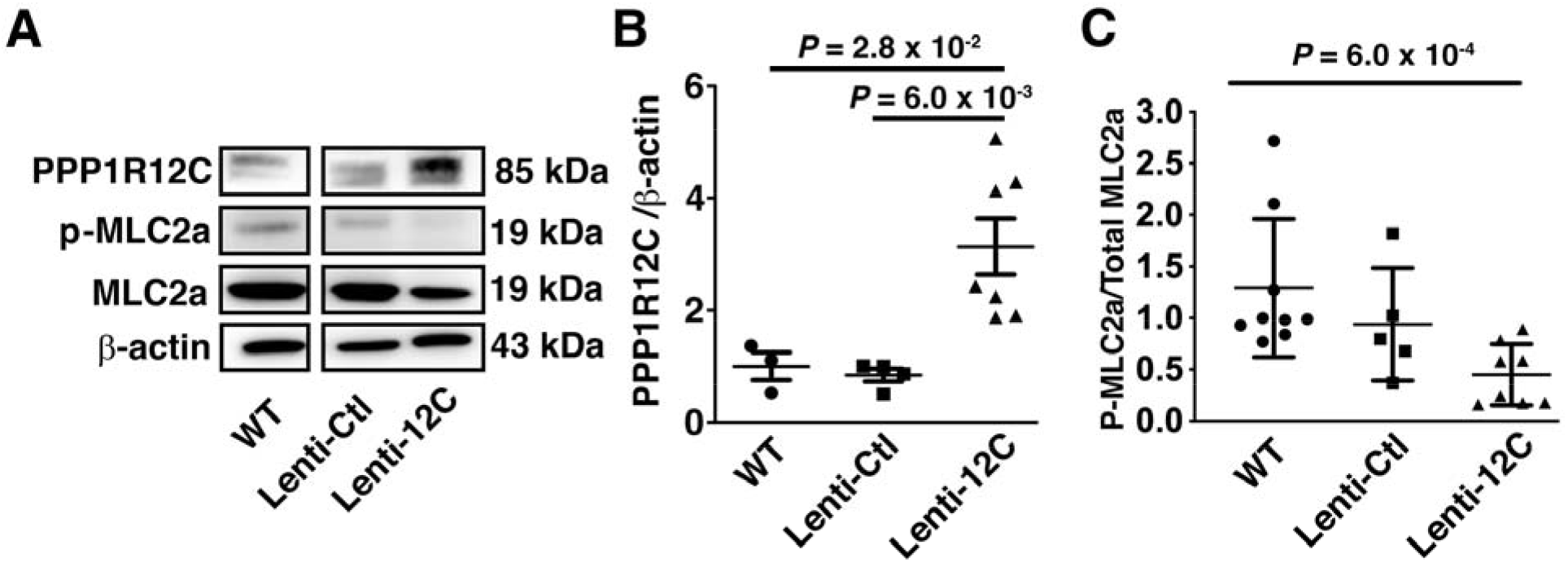
PPP1R12C protein expression is inversely proportional to MLC2a phosphorylation in mice. **A**, Representative Western blot showing increased PPP1R12C abundance in mouse hearts transfected with lentiviral PPP1R12C-GFP construct (Lenti-12C) versus untreated wild-type (WT) and Lentiviral (Lenti-Ctl) controls. **B**, Quantification of PPP1R12C expression in WT versus Lenti-12C mice. n = 3,4,7 in each group. **C**, Quantification of MLC2a phosphorylation among the three mouse models. n = 9,5,8 in each group. Data represent mean±SD. Data for Fig. 4B were determined to have a parametric distribution and an unpaired 2-tail Student’s t-test and ANOVA were used for statistical comparison. Fig. 4C had a non-parametric distribution and data were compared with the Mann-Whitney and Kruskal-Wallis tests. Lenti-12C indicates lentiviral PPP1R12C-treated mice; Lenti-Ctl, lentiviral control mice without PPP1R12C gene insert; MLC2a, atrial myosin light chain 2; PPP1R12C, protein phosphatase 1 regulatory subunit 12C; WT, wild-type control mice.

To evaluate the atrial contractile response to PPP1R12C overexpression, we performed Langendorff perfusion and isolation of atrial cells in the three mouse models, and measured atrial contractile amplitude in response to pacing. We found that atrial contraction amplitude was significantly reduced in Lenti-12C mice versus WT and Lenti- Ctl mice (n = 15,15,15 in each group; *P* = 2.0×10^−4^ and *P* = 1.1×10^−3^, Figure 5A-B), as were contraction duration (*P* = 2.0×10^−6^ vs. WT, Figure 5C), relaxation time (*P* = 7.0×10^−4^ vs. WT, Figure 5D), and time to peak (*P* = 2.9×10^−5^, Figure 5E), an indicator of atrial electrophysiologic latency. There was also a small but significant reduction in sarcomere length in Lenti-12C cells vs. WT (*P* = 1.4×10^−2^, Figure 5F), but no significant differences in peak-to-peak time (*P* = 6.1×10^−1^).

**Figure 5.**
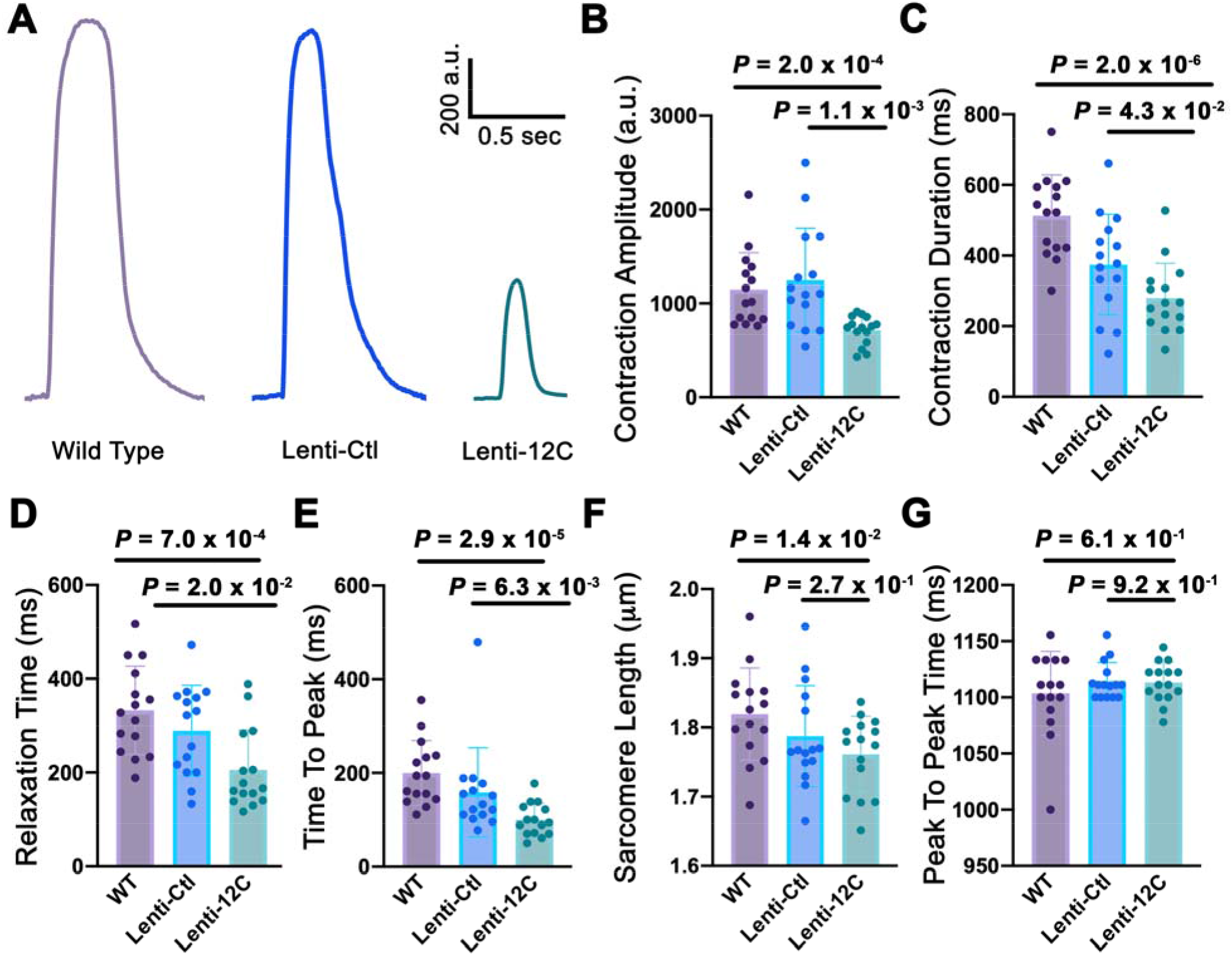
Reduced cell shortening in atrial cells overexpressing PPP1R12C protein. **A**, Representative tracings of contraction amplitude in cells isolated from wild type, Lenti-Ctl, and Lenti-12C mice. **B**, Quantification of contraction amplitude among the three mouse groups. Comparisons of paced cell parameters measured include: **C,** cell contraction duration (ms); **D**, relaxation time (ms); **E**, time to peak amplitude (ms); **F**, sarcomere length (μm); **G**, peak-to-peak time (ms). Data represent mean±SD; comparisons between groups were evaluated with Student’s *t*-test and among groups with one-way ANOVA for data with parametric distribution (Fig. 5C,F), and were evaluated with Mann-Whitney and Kruskal-Wallis tests for data with non-parametric distribution (Fig. 5B, D, E, G). n=15 per group. AU indicates arbitrary units; Lenti-12C, experimental mice treated with lentiviral PPP1R12C insert; Lenti-Ctl, control mice treated with lentivirus without PPP1R12C insert; msec, milliseconds; sec, seconds; WT, wild type mice.

We also evaluated the role of Ca^2+^ handling within Lenti-12C isolated atrial myocytes. We found that in Langendorff-isolated atrial cells, there were no differences in Ca^2+^ release properties including Ca^2+^ transient amplitude (**Supplemental Figure 5**). However, there was a small but significant increase in tau relaxation time constant, which may in part explain the increased latency observed in the cell shortening experiments (**Supplemental Figure 5**). Likewise, we isolated atrial fibers and measured the force-Ca^2+^ relationships among the three groups to determine myofilament response to Ca^2+^ release. We found that mice treated with Lenti-12C have no significant differences in myofilament Ca^2+^-response from Lenti-Ctl and WT controls **(Supplemental Table 3** and **Supplemental Figure 6).** These results are consistent with prior reports of a Ca^2+^ independent pathway for PP1-MLC2 activity due to RhoA activation^34–36^, and indicate that the primary effect of PPP1R12C overexpression is upon myofilament dynamics, and not purely alteration of response to or release of Ca^2+^.

We then sought to determine the functional significance of PPP1R12C expression upon cardiac structural remodeling. Previous investigations of constitutive ablation of MLC2v phosphorylation in mice have shown ultrastructural evidence of: loss of sarcomere organization, irregular Z-lines, non-compact Z-lines, and loss of mitochondria (with mitochondrial remnants) in ventricular cardiomyocytes.^37, 38^ To examine for ultrastructural sarcomeric changes in Lenti-12C mice, we isolated left atrial and left ventricular tissues from Lenti-Ctl and Lenti-12C mice and performed transmission electron microscopy. Similar to previous investigations^37–38^, when we compared left atrial and ventricular tissues from Lenti-Ctl (Figure 6A,C) versus Lenti- 12C mice (Figure 6B,D), we found that Lenti-12C mice demonstrated myofibrillar disarray, altered sarcomere cytostructure, partial or complete loss of the M-line, diffuse/widened Z-line bands, and mitochondrial remnants suggestive of mitochondrial damage. To the best of our knowledge, this is the first evidence of pathologic phosphatase signaling causing myofibrillar disarray at the level of the sarcomere.

**Figure 6.**
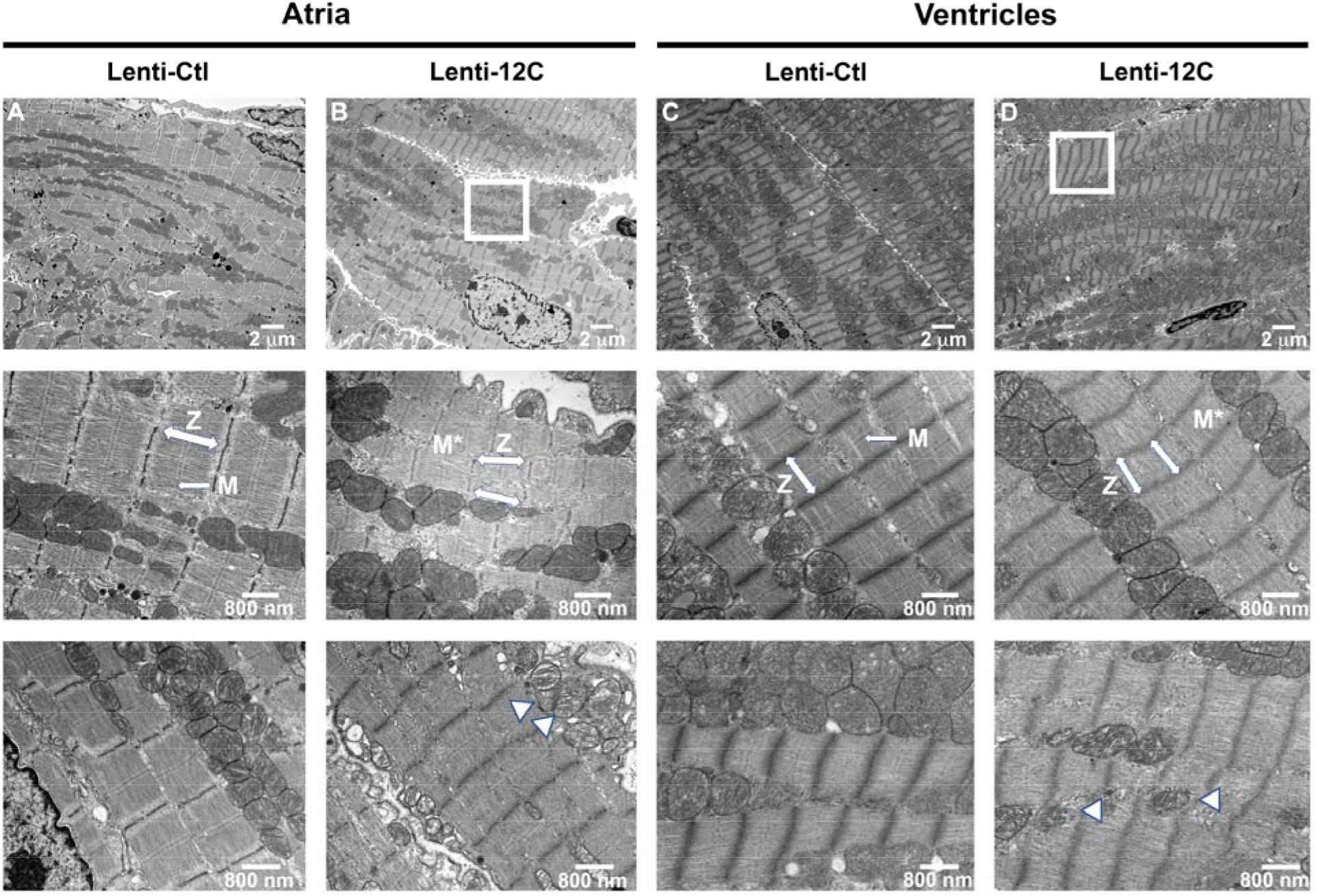
Hypophosphorylation of MLC2a by PPP1R12C causes myofibrillar disarray. **A**, Atria isolated from Lenti-Ctl mice demonstrate neat, ordered sarcomeres (top, 2 μm), distinct Z-line and M-line structures (middle, 800 nm), and normal mitochondria (bottom, 800 nm). **B**, Atria from Lenti-12C mice demonstrate myofibrillar disarray and altered sarcomere cytostructure (box, top, 2 μm), partial or complete loss of M-line (M*) and indistinct Z-line bands (middle, 800 nm), and evidence of mitochondrial membrane damage (arrowheads, bottom, 800 nm). **C**, Ventricles isolated from Lenti-Ctl mice demonstrate neat, ordered sarcomeres as described in (**A**). **D**, Ventricles from Lenti-12C mice demonstrate myofibrillar disarray and altered sarcomere cytostructure (box, top, 2 μm), partial or complete loss of M-line (M*) and indistinct Z-line bands (middle, 800 nm), and evidence of mitochondrial membrane damage (arrowheads, bottom, 800 nm).

Next, we examined Lenti-12C mice for evidence of atrial and ventricular remodeling using high frequency echocardiography. We found that at 2 months of age, Lenti-12C mice had a significant increase in atrial diameter versus Lenti-Ctl and WT controls (n = 12,8,12 in each group; *P* = 5.0 × 10^−6^; Figure 7A). Follow-up studies focused on atrial function, in collaboration with Dr. Goergen’s group, showed a reduction in both atrial ejection fraction (n = 5,5,5 in each group; *P* = 3.0×10^−3^ vs. WT*; P* = 1.1×10^−2^ vs. Lenti-Ctl; Figure 7B-D) and atrial strain (n = 5,5,5 in each group; *P* = 4.0×10^−3^ vs. WT*; P* = 2.0×10^−3^ vs. Lenti-Ctl; Figure 7E) in Lenti-12C mice versus WT and Lenti-Ctl mice. There was also a significant increase in left ventricular (LV) posterior wall thickness in diastole (LVPWd) and end-diastolic diameter (EDD) versus controls, signifying mild ventricular dilation, and which is commonly associated with AF in human patients (Table 1, **Supplemental Table 4**).^39, 40^ However, there was no significant difference in left ventricular ejection fractions (LVEF) among the three groups (Figure 7F). Similarly, we compared the relationship between atrial and ventricular ejection fractions (EF) in the three models and found no significant correlation among the three groups (Figure 7G).

**Figure 7.**
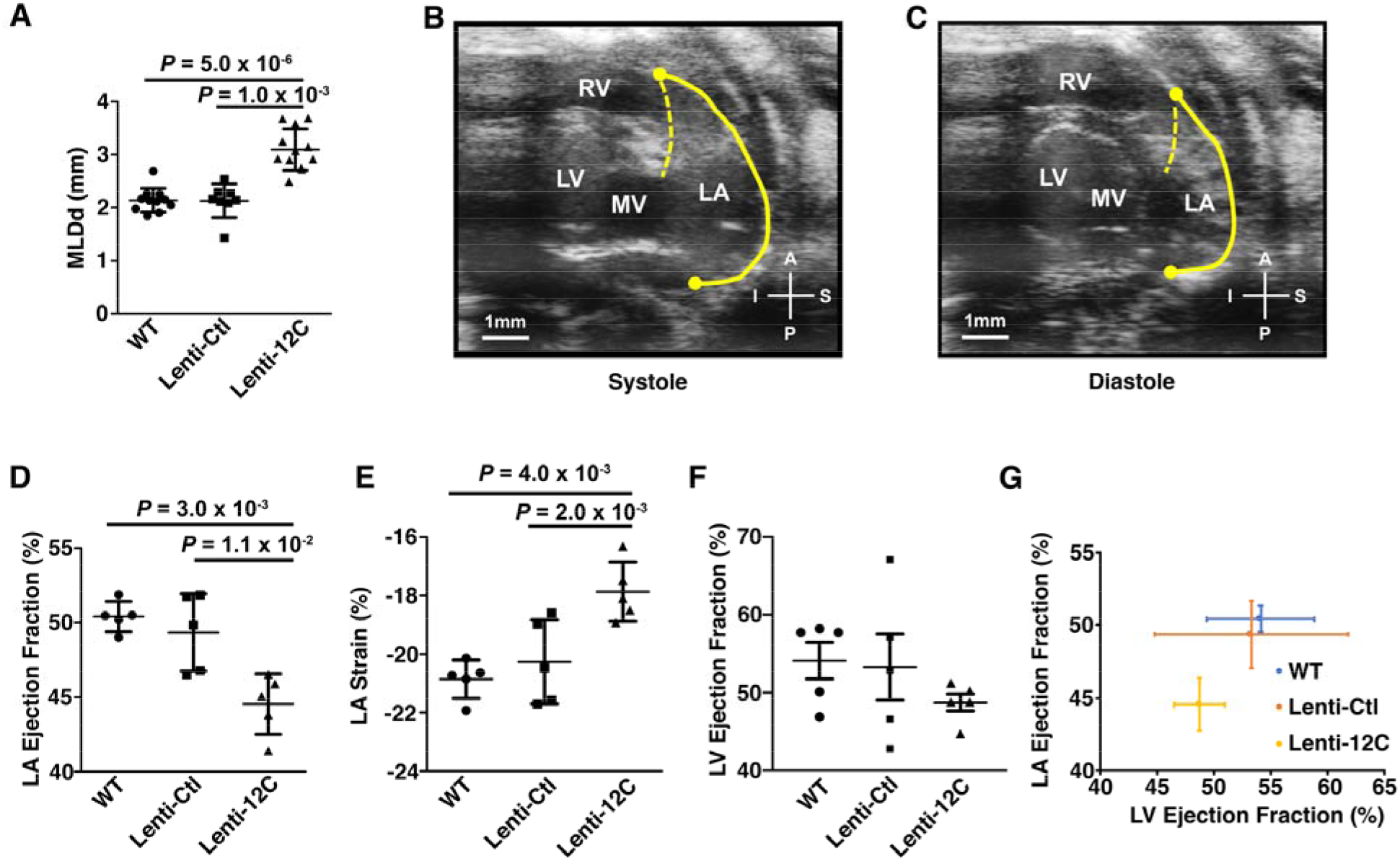
Increased PPP1R12C expression causes atrial enlargement and atrial hypocontractility. **A**, Echocardiographic measurements of medio-lateral diameter of the left atrium during diastole. **B-C**, Representative echocardiogram images of the left atrial walls during systole (**B**) and diastole (**C**). **D**, Quantification of left atrial ejection fraction. **E**, Quantification of left atrial strain. **F**, Quantification of left ventricular ejection fraction. **G**, Plot of left atrial ejection fraction (y-axis) versus left ventricular ejection fraction (x-axis). Data are plotted with error bars indicating SD; the Shapiro-Wilk test determined parametric distribution of the data and comparisons among groups were evaluated with Student’s *t*-test and one-way ANOVA. n = 12,8,12 for panels A-C; n = 5,5,5 in each group for panels D-G. MLDd indicates medio-lateral diameter of the left atrium during diastole; PPP1R12C, protein phosphatase 1 regulatory subunit 12C; Lenti-12C, lentiviral PPP1R12C-treated mice; Lenti-Ctl, lentiviral control mice without PPP1R12C gene insert; WT, wild-type control mice.

**Table 1.**
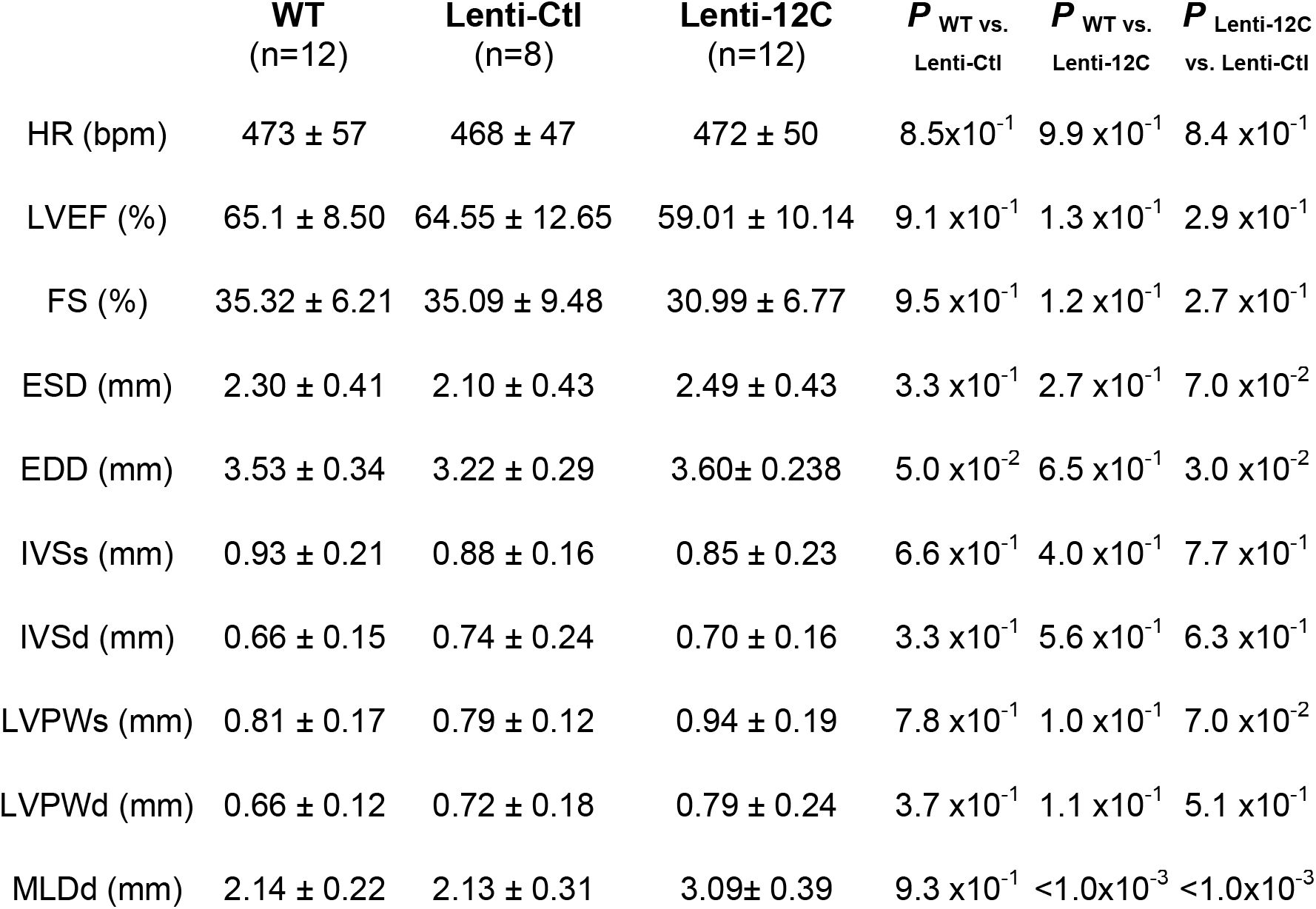
Comparison of common atrial and ventricular parameters on echocardiography among the three mouse groups. Data distribution was determined to be parametric using the Shapiro-Wilk test, and were compared using Student’s *t*-test and ANOVA. EDD indicates end diastolic diameter of the left ventricle; ESD, end systolic diameter left ventricle; FS, fractional shortening; HR, heart rate; IVSd, interventricular septum thickness in end-diastole; IVSs, interventricular septum thickness in end-systole; Lenti-12C, lentiviral PPP1R12C-treated mice; Lenti-Ctl, lentiviral control mice without PPP1R12C gene insert; LVEF, left ventricular ejection fraction; LVPWd, left ventricular posterior wall dimension in end-diastole; LVPWs, left ventricular posterior wall dimension in end-systole; MLDd, medio-lateral diameter of the left atrium during diastole; PPP1R12C, protein phosphatase 1 regulatory subunit 12C; WT, wild-type control mice.

| | WT<br>(n=12) | Lenti-Ctl<br>(n=8) | Lenti-12C<br>(n=12) | $P_{WT \text{ vs. } Lenti-Ctl}$ | $P_{WT \text{ vs. } Lenti-12C}$ | $P_{Lenti-12C \text{ vs. } Lenti-Ctl}$ |
| --- | --- | --- | --- | --- | --- | --- |
| HR (bpm) | 473 ± 57 | 468 ± 47 | 472 ± 50 | 8.5x10<sup>-1</sup> | 9.9 x10<sup>-1</sup> | 8.4 x10<sup>-1</sup> |
| LVEF (%) | 65.1 ± 8.50 | 64.55 ± 12.65 | 59.01 ± 10.14 | 9.1 x10<sup>-1</sup> | 1.3 x10<sup>-1</sup> | 2.9 x10<sup>-1</sup> |
| FS (%) | 35.32 ± 6.21 | 35.09 ± 9.48 | 30.99 ± 6.77 | 9.5 x10<sup>-1</sup> | 1.2 x10<sup>-1</sup> | 2.7 x10<sup>-1</sup> |
| ESD (mm) | 2.30 ± 0.41 | 2.10 ± 0.43 | 2.49 ± 0.43 | 3.3 x10<sup>-1</sup> | 2.7 x10<sup>-1</sup> | 7.0 x10<sup>-2</sup> |
| EDD (mm) | 3.53 ± 0.34 | 3.22 ± 0.29 | 3.60 ± 0.238 | 5.0 x10<sup>-2</sup> | 6.5 x10<sup>-1</sup> | 3.0 x10<sup>-2</sup> |
| IVSs (mm) | 0.93 ± 0.21 | 0.88 ± 0.16 | 0.85 ± 0.23 | 6.6 x10<sup>-1</sup> | 4.0 x10<sup>-1</sup> | 7.7 x10<sup>-1</sup> |
| IVSd (mm) | 0.66 ± 0.15 | 0.74 ± 0.24 | 0.70 ± 0.16 | 3.3 x10<sup>-1</sup> | 5.6 x10<sup>-1</sup> | 6.3 x10<sup>-1</sup> |
| LVPWs (mm) | 0.81 ± 0.17 | 0.79 ± 0.12 | 0.94 ± 0.19 | 7.8 x10<sup>-1</sup> | 1.0 x10<sup>-1</sup> | 7.0 x10<sup>-2</sup> |
| LVPWd (mm) | 0.66 ± 0.12 | 0.72 ± 0.18 | 0.79 ± 0.24 | 3.7 x10<sup>-1</sup> | 1.1 x10<sup>-1</sup> | 5.1 x10<sup>-1</sup> |
| MLDd (mm) | 2.14 ± 0.22 | 2.13 ± 0.31 | 3.09 ± 0.39 | 9.3 x10<sup>-1</sup> | <1.0x10<sup>-3</sup> | <1.0x10<sup>-3</sup> |

Finally, we sought to determine whether PPP1R12C overexpression contributes to AF risk. At a young 2-3 month age, we found no AF at baseline in each of the three groups. However, when we performed trans-esophageal atrial pacing with burst and extrastimulus protocols (**see Methods**^41^), we found a significant increase in AF burden in Lenti-12C mice versus both Lenti-Ctl and WT controls (n = 6,6,5 in each group; *P* = 1.8×10^−2^ vs. WT and *P* = 4.1×10^−2^ vs. Lenti-Ctl, Figure 8A-B). These data demonstrate that PPP1R12C overexpression not only contributes to atrial hypocontractility in AF, but is a risk factor for the initiation of AF as well.

**Figure 8.**
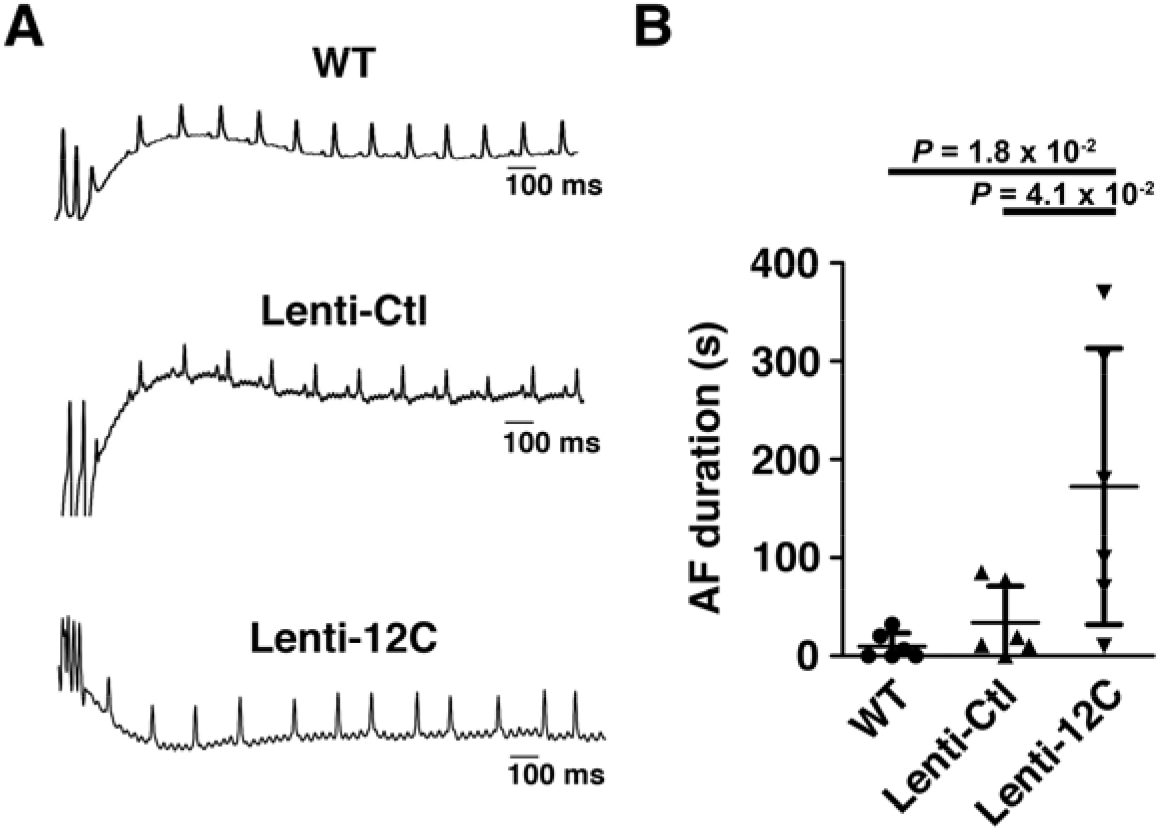
Overexpression of PPP1R12C increases pacing-induced atrial fibrillation. **A**, Representative surface electrocardiograms in mice receiving trans- esophageal atrial pacing. Lenti-12C mice are prone to pacing-induced AF. **B**, Quantification of AF burden among the four groups. Data are plotted with error bars indicating SD. Data were determined to have a parametric distribution by the Shapiro-Wilk test, and Student’s *t*-test was used to compare differences between groups and one-way ANOVA was used among groups. AF indicates atrial fibrillation; PPP1R12C, protein phosphatase 1 regulatory subunit 12C; Lenti-12C, lentiviral PPP1R12C-treated mice; Lenti-Ctl, lentiviral control mice without PPP1R12C gene insert; WT indicates wild-type control mice.

## DISCUSSION

### PPP1R12C Promotes MLC2a Dephosphorylation and Atrial Hypocontractility

In these studies, we show for the first time that: 1) human AF patients demonstrate significantly increased atrial expression of PPP1R12C, 2) PPP1R12C is a major regulator of MLC2a phosphorylation in the heart, 3) PPP1R12C activity is modifiable at the MRCK site and effects changes in MLC2a phosphorylation, and 4) PPP1R12C overexpression contributes, in part, to AF susceptibility *in vivo*. Our studies are among the first to provide mechanistic insights into atrial hypocontractility in AF, and complement previous observations of myofilament signaling in AF.^13–16^ Previous studies of PPP1R12C function in neuronal tissues suggest that PPP1R12C is a regulatory scaffold upon which PP1c dephosphorylates MLC2.^19^ However, the expression and physiologic role of PPP1R12C in the heart had remained unstudied. Given that MLC2 phosphorylation is a major regulator of cardiac contractility, and recent evidence of altered PP1 R-subunit expression in human AF, our studies are also the first to selectively examine the role of PP1 regulatory proteins upon the PP1-MLC2a interface.^42^

### Molecular Insights into Atrial Hypocontractility in AF

To date, there are two complementary strategies that have been used to evaluate atrial remodeling in AF: 1) evaluation of tissues derived from AF patients and/or animal models of tachypacing-induced AF to determine contributory molecular mechanisms, and 2) evaluation of specific pathways implicated in AF to determine contribution to the AF phenotype. The first approach has yielded valuable insights into the global AF-related cell and tissue level remodeling seen in AF. El-Armouche *et al*. showed that human patients with chronic AF have a complex atrial cellular phenotype.^13^ Markers of atrial remodeling included a higher ratio of phospholamban to SERCA2a, higher PP1 and PP2A activities, lower phosphorylation of myosin binding protein C, and enhanced protein kinase A phosphorylation of phospholamban; however, the degree of MLC2a phosphorylation was unknown because it was not detected in this study. Likewise, animal models have given us valuable insight into the progression of atrial tachyarrhythmias and their effects upon atrial remodeling. Greiser *et al*. compared goats with AV-block/pacing induced atrial dilation versus an atrial repetitive burst pacing AF model.^14^ This seminal work showed that atrial burst pacing had a larger reduction in atrial work index, reduced force of contractility, reduced action potential, and reduced phosphorylation of phospholamban, associated with reduced SR Ca^2+^ load, and suggested that atrial remodeling in AF exhibits unique molecular patterns distinct from pressure-overload related atrial remodeling.^14^

In a clinically relevant model of atrial remodeling in heart failure, Yeh *et al*. demonstrated that in a dog model of ventricular tachypacing, Ca^2+^ handling is a common mechanism uniting both ventricular and atrial dysfunction.^43^ Atrial cell shortening in this model was reduced by 50%, and was associated with reduced MLC2a phosphorylation and myosin binding protein C phosphorylation, whereas troponin I expression and phosphorylation were similar to controls. These atrial remodeling findings were further confirmed by the Nattel group in a 7 day atrial tachypacing trial in dogs versus sham (unpaced) control.^16^ This work laid an important foundation for the current study, as it was the first example of reduced MLC2a phosphorylation in AF, and suggested a major role for MLC2a phosphorylation in atrial contractile dysfunction.

These investigations have been valuable for several reasons. They have facilitated the comparison of different atrial pathologies from a variety of pathological inputs, and they have allowed a systematic translational approach to understanding multiple concurrent molecular pathways. However, AF is a complex cellular phenotype and these approaches were not designed to dissect specific pathways. Additionally, among animal models, atrial pacing may not fully recapitulate the complex AF phenotype experienced in AF-prone human patients. Therefore, the complementary approach to specific pathway analysis has been useful to further examine molecular contributions from specific pathways.

Second, genetic animal models have been useful to dissect specific molecular pathways contributing to AF. We and others have previously used genetic models to detail how alterations in protein kinase activity have led to pathologic alterations in Ca^2+^ release ^28, 44^, ion channel dysfunction ^45, 46^, and myofilament signaling in AF.^47^ However, an underappreciated but growing area of study is the recognition that protein phosphatases underpin a significant degree of pathologic signaling in AF. Importantly, protein phosphatase specificity is distinct from protein kinases, in that the regulatory subunits define subcellular site specificity. This is especially true for PP1, the most abundant cardiac protein phosphatase.^48^ As Chiang *et al*. previously noted: “Because there are only a few different PP1c isoforms, all of which share a high degree of homology, the spatial and temporal specificity of PP1 for different targets is largely regulated by association with these R-subunits.”^17^ Chiang *et al*. went on to demonstrate the relevance of PPP1R in human AF, showing that PPP1R7 may act as a “sponge” for the catalytic PP1c subunit, and can alter its subcellular localization during atrial remodeling. Alsina *et al*. found that PPP1R3A, a PP1 R-subunit targeting both phospholamban and the cardiac ryanodine receptor, protein expression is downregulated in human patients with AF, and that genetic ablation of *Ppp1r3a* in mice results in impaired binding of PP1 to both Ca^2+^ regulating proteins.^42^ Thus, there is a growing body of evidence that PP1 R-subunits play a major role in atrial remodeling.

PPP1R12C, a previously understudied PP1 subunit, was recently recognized as a major regulator of myosin and actin dynamics, and which participates in cytoskeleton rearrangements necessary for effective antiviral responses.^49^ In the heart, PPP1R12C was previously found to be overexpressed in atrial tissues of patients with AF (versus sinus rhythm), however the functional significance of this finding was unclear.

In our study, we sought to evaluate PP1 regulation of a major determinant of atrial contractility, MLC2a. As PPP1R12C activity in the heart had not previously been described, we began with *in vivo* and *in vitro* experiments to elucidate the function of PPP1R12C in the heart. We found that increased PPP1R12C expression is associated with increased PPP1R12C-PP1c binding and targeting of PP1c to sarcomeric MLC2a, resulting in enhanced dephosphorylation of MLC2a. We determined that PPP1R12C protein activity is regulated at the MRCK site, and this site is a pharmacologic target, as evidenced by exposure to BDP5290, an experimental chemotherapeutic agent which enhances PPP1R12C-PP1c binding and activity (Figure 3). Future pharmacologic investigations will be necessary to discover an inhibitor of PPP1R12C, which is expected to increase MLC2a phosphorylation, increase atrial contractility, and reduce incidence of both AF and stroke.

We also found an indispensable role for PPP1R12C in regulating ventricular contractility. Lenti-12C mice developed a mild dilated cardiomyopathy phenotype, highlighting the negative effects of unchecked MLC2 dephosphorylation, and further suggested that homeostasis of MLC2-P is a finely titrated event in myocardial signaling. Upcoming studies will examine the relevance of PPP1R12C activity in human heart failure. Finally, we demonstrated that overexpression of PPP1R12C contributes to AF risk *in vivo*. While the mechanisms of arrhythmogenesis in AF are complex, atrial contractility has previously been shown to affect AF risk primarily through alterations in phosphorylation of sarcomeric proteins^50^, and which was present via MLC2a dephosphorylation in our Lenti-12C mice.

### Clinical Insights into Atrial Contractile Dysfunction

Our findings have important clinical implications for patients with AF. Atrial dilation and related hypocontractility are major risk factors for stroke, and also AF recurrence. Currently, the only limb of Virchow’s triad of vascular thrombosis unaddressed for stroke risk in AF patients is atrial hypocontractility. Atrial hypocontractility, a term often used interchangeably with atrial stunning, is well described in echocardiography where spontaneous echo contrast fills the left atrium and LAA during AF.^51^ Detailed echocardiographic measurements of mitral inflow, pulmonary venous flow, and atrial wall contractility in human AF patients show there is a significant reduction in atrial contractility in AF.^52, 53^ Interestingly, after either electrical or pharmacologic cardioversion, there is a persistent reduction in LA contractility and spontaneous echo contrast that may persist up to 6-8 weeks.^54^ The observation that atrial hypocontractility persists after conversion from AF to normal sinus rhythm suggests that cellular remodeling in AF may create a substrate for atrial hypocontractility. By altering this hypocontractile atrial substrate, reducing PPP1R12C expression may not only prevent adverse atrial remodeling, but also increase atrial contractile function and reduce stroke risk in AF-susceptible patients.

### Limitations

We acknowledge that while these investigations have shed light on the roles of PPP1R12C in AF, our study does have limitations. The mechanisms underlying AF are complex^16^, and we have selectively examined one important molecular aspect of hypocontractility in AF, the PP1-myofilament interface. However, our studies show that the expression and phosphorylation of other relevant myofilament proteins are unaltered with overexpression of PPP1R12C in mice, suggesting that the PPP1R12C-MLC2a interface is a critical regulatory pathway for atrial contractility. Second, there are inherent limitations in using viral vectors in mice to model human disease; we utilized this model to be able to finely titrate PPP1R12C expression close to that experienced in human AF patients. However, this mouse model may not fully recapitulate the AF phenotype observed in humans, and the viral vector may mildly reduce cardiac contractility itself. Similarly, mice do not typically develop stroke in the condition of atrial fibrillation. Third, we utilized an alpha-MHC promoter in our lentiviral vector, which promotes both atrial and ventricular expression of PPP1R12C. Ni et al. have recently described an atrial-specific gene delivery system using an adeno-associated viral vector and atrial natriuretic factor (ANF) promoter; in the future, lentiviral approaches atrial-specific gene delivery may allow better atrial selectivity.^55^

Additionally, this study primarily evaluates global MLC2a phosphorylation, however, gradients between the epi- and endocardium have been shown in the ventricle to be important to adjust the function to the stress^38^; this might also apply to the atrium as well. Finally, with regards to echocardiographic techniques used, all metrics were computed based on two-dimensional views of the heart and thus relied on the sonographer to consistently reproduce similar image planes between mice.^56, 57^ To reduce observer bias, we utilized two separate echocardiography readers at two institutions (UIC and Purdue) and blinded the readers to the treatment modality. Despite these limitations, we show for the first time that PPP1R12C plays a major mechanistic role for atrial hypocontractility in AF. Future studies are needed to determine whether selective blockade of the PP1c-PPP1R12C interaction may increase MLC2a phosphorylation and prevent atrial stunning and stroke risk in AF-susceptible individuals.

## NON-STANDARD ABBREVIATIONS AND ACRONYMS

AF: Atrial fibrillation
BDP: Drug BDP5290
cAF: Chronic atrial fibrillation
DMSO: Dimethyl sulfoxide
EP: Electrophysiology studies, invasive
Lenti-12C: Mice treated with lentiviral PPP1R12C vector
Lenti-Ctl: Mice treated with lentiviral GFP vector control, without PPP1R12C insert
LV: Left ventricle or left ventricular
MLC2a: Atrial myosin light chain 2
MRCK: Myotonic dystrophy kinase-related Cdc-42 binding kinase
PP1c: Protein phosphatase 1 catalytic subunit
PPP1R12C: Protein phosphatase 1 regulatory subunit 12C
RhoA: Ras homolog family member A
SERCA: Sarco/endoplasmic reticulum Ca^2+^ ATPase
SR: Sinus rhythm

## ACKNOWLEDGMENTS

M.D.M. would like to thank Mr. Louis Mange for his friendship, mentorship, and financial support that made this project possible. We also thank Ramona Löcker, Simone Olesch and Annette Kötting-Dorsch for expert technical assistance.

